# Prawn aquaculture as a method for schistosomiasis control and poverty alleviation: a win-win approach to address a critical infectious disease of poverty

**DOI:** 10.1101/465195

**Authors:** Christopher M. Hoover, Susanne H. Sokolow, Jonas Kemp, James N. Sanchirico, Andrea J. Lund, Isabel Jones, Tyler Higginson, Gilles Riveau, Amit Savaya-Alkalay, Shawn Coyle, Chelsea L. Wood, Fiorenza Micheli, Renato Casagrandi, Lorenzo Mari, Marino Gatto, Andrea Rinaldo, Javier Perez-Saez, Jason R. Rohr, Amir Sagi, Justin V. Remais, Giulio A. De Leo

**Author notes:** Now at Google Brain (Jonas Kemp). Denotes shared senior authorship (Justin V. Remais, Giulio A. De Leo).

## Abstract

Recent evidence suggests crustacean snail predators may aid schistosomiasis control programs by targeting the environmental component of the parasite’s life cycle through predation of the snail species that serve as intermediate hosts of the parasite. We evaluate costs, benefits, and potential synergies between schistosomiasis control and aquaculture of giant prawns using an integrated bio-economic-epidemiologic model. We identified combinations of stocking density and aquaculture cycle length that maximize profit and offer disease control benefits for sustainable schistosomiasis control. We consider two prawn species in sub-Saharan Africa: the endemic, non-domesticated *Macrobrachium vollenhovenii*, and the non-native, domesticated *Macrobrachium rosenbergii*. We find that, at profit-optimal densities, both *M. rosenbergii* and *M. vollenhovenii* can complement conventional control approaches (mass drug treatment of people) and lead to sustainable schistosomiasis control. We conclude that integrated aquaculture strategies can be a win-win strategy in terms of health and sustainable development in schistosomiasis endemic regions of the world.

---

Schistosomiasis is a debilitating disease of poverty, affecting around 200 million people worldwide ^1,2^. It is caused by trematode parasites of the genus *Schistosoma* that undergo a life cycle involving passage between definitive human hosts and freshwater snails that act as intermediate hosts. While safe and effective treatments, such as the anthelmintic drug Praziquantel, are available to reduce parasite burden and associated symptoms from infected individuals, rapid reinfection in highly endemic areas leads to persistent hotspots of infection^3,4^. Successful long-term elimination efforts may require strategies that go beyond conventional mass drug administration (MDA) campaigns to explicitly target the environmental reservoir of the disease ^5^.

One option for reducing transmission is cultivating snail predators, such as river prawns, via aquaculture. River prawns have been shown to reduce schistosomiasis transmission by consuming snails in the aquatic environment where people contact infested water ^14,15^. Given that schistosomiasis is a disease of poverty, combining the nutritional and economic benefits of prawn aquaculture with disease control via prawn predation on snails may offer a sustainable approach to combat schistosomiasis and improve well-being and economic development in endemic areas. Here, we develop an integrated bio-economic-epidemiologic model to investigate whether extensive prawn aquaculture (using either endemic *M. vollenhovenii* or nonbreeding monosex aquaculture of *M. rosenbergii*) can be managed at schistosome transmission sites such as rice paddies or enclosed points of water contact where people are exposed ^15^ to simultaneously maximize profit and control schistosomiasis.

There is a rich history of environmental interventions for schistosomiasis that target the intermediate snail hosts ^6^. Molluscicides are effective in reducing snail populations and have been used in integrated campaigns to control schistosomiasis in areas of South America, Northern Africa, and Southeast Asia ^6–8^. However, these approaches generally require repeated applications of chemicals that may negatively affect non-target species in addition to *Schistosoma*-bearing snails ^9,10^. A more sustainable approach for snail control involves the introduction of snail predators, such as river prawns, which have been shown to actively seek out and consume important intermediate hosts for human schistosomes in laboratory settings, including *Biomphalaria* and *Bulinus* snails ^11–13^. Field trials in which crustacean predators have been introduced to reduce schistosomiasis transmission have been successful in reducing reinfection rates in humans following MDA ^14,15^.

In addition to being voracious predators of snails ^13,16^, river prawns are a valuable food commodity^17,18^. The giant freshwater prawn *Macrobrachium rosenbergii* has been domesticated and widely used in commercial hatchery-based aquaculture ^19^, providing both a key source of protein and encouraging local economic development ^20^. Furthermore, advances in the production of nonbreeding *M. rosenbergii* monosex populations reduces the risk of prawn invasion in areas this species is not native, suggesting safe use of this biological control agent globally ^21,22^. In sub-Saharan Africa, where at least 90% of schistosomiasis cases occur ^1,23^, the native African river prawn *M. vollenhovenii* has been proposed as an alternative to *M. rosenbergii* for aquaculture ^21^. Research into scalable *M. vollenhovenii* aquaculture is ongoing, as the use of this native species may be more attractive due to its historical presence in the local river ecology in western Africa.

Extensive prawn aquaculture, increasingly common in developing countries ^17–19^, consists of large enclosures, low prawn densities, and no use of supplemental feed, substrate, or additional oxygenation. As such, extensive aquaculture is more compatible with the water resource management needs of rural communities in key areas of sub-Saharan Africa, and can be easily integrated into rice agriculture that is increasingly an important part of food production and is present in many schistosomiasis endemic areas ^24,25^.

Scaling the prawn aquaculture approach for schistosomiasis control requires that the economic goals of prawn aquaculture are compatible with the public health goal of schistosomiasis control. Thus, we seek to identify conditions under which schistosomiasis reinfection in the human population may be curbed while maximizing the economic benefit of prawn aquaculture. We first develop a bio-economic production model of *Macrobrachium* spp. aquaculture to identify the stocking density and duration of the grow out phase that maximizes profit for *M. rosenbergii* and *M. vollenhovenii*. Then, we expand an epidemiologic model of schistosomiasis transmission dynamics to include snail size structure and infection dynamics. This model is coupled with the aquaculture model through size- and density-dependent prawn predation, and parameterized via results of previous laboratory and field-based empirical studies. We use estimates of the disability adjusted life years (DALYs) lost due to schistosomiasis infection derived from the integrated model to compare aquaculture-based prawn interventions with conventional MDA interventions, and to estimate the benefits of utilizing both MDA and prawn aquaculture for schistosomiasis control. We conclude with an extensive sensitivity analysis to evaluate the feasibility of such interventions in a variety of settings.

## Methods

The integrated model has three components: a) a bio-economic aquaculture component, simulating yields and accounting for density-dependent mortality and somatic growth of *Macrobrachium* spp. prawns over a range of initial stocking densities; b) an epidemiologic component to simulate the dynamics of mean schistosome worm burden in humans and population and infection dynamics of intermediate-host snails through a size structured, SEI compartmental model; and c) a predation function describing the rate at which prawns consume snails as a function of snail density and of snail and prawn body sizes, which links the epidemiologic and aquaculture models.

### (a) Aquaculture model

We assume that a necessary supply chain of hatcheries and nurseries supplies juvenile prawns to stock at the desired transmission sites. The dynamics of a cohort of *P*_0_ juvenile prawns of initial mean length, *L*_0_ [mm], stocked at time *t* = 0 *days* in an enclosure of 1,000 *m*^2^—a size consistent with either a large water contact site or a typical rice field in small-scale, subsistence agriculture settings—are simulated over time as a function of density-dependent and size-dependent growth and mortality.

Adult *Macrobrachium rosenbergii* males can be grouped in three different categories that grow at different rates depending upon their body size and developmental phase: small males (SM) between 5-20g, orange-clawed males (OC) between 30-180g, and blue-clawed male (BC) reaching up to 250 g, with growth of some smaller prawns being suppressed by the largest BC males ^26^. Though this same social structure has not been investigated for *M. volenhovenii*, we assume here that it is the same. Growth of individual crustaceans is typically stepwise and occurs through a sequence of molting events, but here the population-average growth of prawns is modeled as somatic growth with the von Bertalanffy growth equation (VBGE) ^27^:

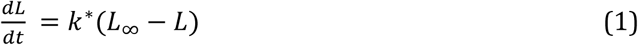

VBGE produces a classic increasing-and-saturating growth curve with length *L* at time *t* (days after stocking) eventually approaching the mean asymptotic length, *L_∞_*. Based on experimental stocking trials showing reduced growth rates of *M. rosenbergii* at high stocking densities ^26^, a modified Brody growth coefficient, *k*^∗^, was estimated as a decreasing function of total population biomass, *Ω*:

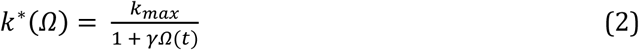

where *k_max_* is the maximum value of the Brody coefficient at low densities and *γ* a coefficient parameterized to produce a density-dependent reduction in somatic growth qualitatively resembling that observed in experimental trials ^26^.

Total population biomass, *Ω*(*t*), is computed as the product of mean prawn body size in grams, *B*(*t*), and population size, *P*(*t*), *t* days after stocking:

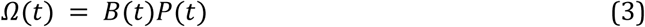

Body size *B*(*t*) is derived as an allometric function of prawn length, *L*(*t*), from *M. rosenbergii* grow-out data ^28,29^:

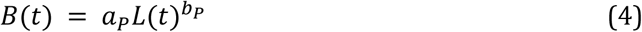

As prawns are generalist consumers with a wide range of invertebrate fauna and organic detritus in their diet ^30^, growth as described by eq. 1-4 is assumed independent from snails’ density and the corresponding predation rates.

After stocking, the total number of prawns in the enclosure decreases over time, with *per*-*capita* mortality rate modeled as an additive function of two components: (*i*) an exponentially decreasing function of body size, *B*(*t*), as large prawns exhibit lower mortality than small prawns ^31^; (ii) a linearly increasing function of total population biomass, Ω(*t*), which accounts for density-dependent competition for resources and cannibalism at high population densities ^26^. Accordingly, the dynamics of a cohort of initial size *P*_0_ was described as follows:

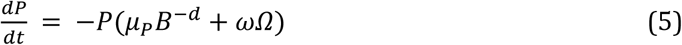

with parameters *μ_P_* and *d* derived from previous studies ^31,32^ and *ω* parameterized to produce density-dependent mortality outcomes qualitatively similar to those observed in the experimental trials by Ranjeet and Kurup ^26^. Natural recruitment is excluded from the aquaculture model, as new prawns enter the system only in discrete, exogenously controlled events, when *P*_0_ prawns of initial average body size *L*_0_ are stocked from nurseries and grown out to market size. We assume natural, size- and density-dependent mortality are the only causes of prawn population decline and do not consider other sources such as predation by e.g. seabirds, escape from enclosures or rice fields, disease outbreaks, or declines in water quality that may affect prawn health.

Prawns weighing <30g are generally not of commercial interest, therefore only OC and BC males are considered marketable. Experimental trials by Ranjeet and Kurup with *M. rosenbergii* ^26^ showed the fraction of retrievable, market-size (>30g) prawns decreases linearly with increasing prawn stocking density. Accordingly, commercial yield at the end of a production cycle of length *t* = *T* is only a fraction of the total biomass:

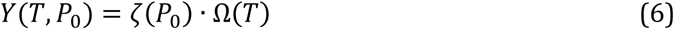

where *ζ*, the fraction of marketable size prawns in the population, is a decreasing function of initial stocking density *P*_0_ estimated from data in the Ranjeet and Kurup experiments ^26^.

Cumulative profits over a finite time horizon are determined by the profit produced per cycle and the number of cycles completed within the given time period. In a time period of *T_max_* days, the number of aquaculture cycles completed for each *Macrobrachium* species (sp) is 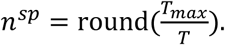 Cumulative discounted profit for each species, *CP^sp^*, is then estimated as the sum of net discounted revenue for every cycle completed by *T_max_*:

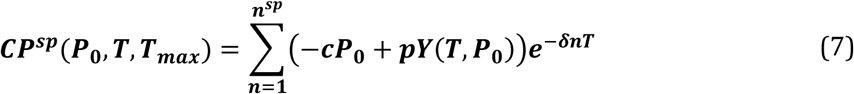

where *p* is the price per unit weight (USD/kg), *c* the per capita cost of stocked juvenile prawns, and *δ* the discount rate to account for the time lag between initial stocking costs and delayed revenues of commercial size prawns. Following the comprehensive price analysis by Dasgupta and Tidwell, we set *c* = $0.10/*P* for a juvenile prawn of *L*_0_ = 40*mm* (corresponding to ~0.35*g* juvenile prawns) and *p* = $12/*kg* harvested ^33^. The discount rate is set to 7%, which is likely on the low end for sub-Saharan African countries endemic with schistosomiasis but higher than the 3-4% rate used for discounting long term government projects in the United States ^34^. Other costs such as maintaining nurseries to produce juvenile prawns are considered external to the aquaculture scenario considered here and are therefore not included in the profit estimation. The influence of such costs are considered in additional sensitivity analyses described below.

Cumulative discounted profits are maximized by jointly choosing the rotation length, *T*, and the initial stocking density, *P*_0_. Given the rotation length, the number of rotations in the time period is determined, as *T_max_* is given. Equation 7 has it basis in the optimal rotation models in forestry (see, e.g., ^35^) and commercial aquaculture operations (see, e.g., ^36,37^) which balance the marginal benefits of further growth against the opportunity costs of waiting to harvest. The resulting optimal rotation length is therefore shorter than a simple rule of when to harvest based solely on maximizing growth dynamics. In our setting, we expect the same trade-off between benefits from waiting to harvest the prawns at a larger size against the opportunity costs of delaying the economic returns from future harvests.

Parameters used in the prawn aquaculture model simulations are listed in Table S1.

### (b) Epidemiologic model

Building on our previous modeling of *S. haematobium* ^15,38^, the infection dynamics of the intermediate host snail population, ***N***, are modeled as snails transition between *i* ∈ {*S*, *E*, *I*} infection compartments corresponding to susceptible (*S*), exposed (*E*, pre-patent), and infected (*I*, patent) states. Furthermore, the growth dynamics of snails are modeled as they move through *j* ∈ {1, 2, 3} size classes corresponding to 4*mm*, 8*mm*, and 12*mm* mean snail diameter (Fig. 1). The model is further extended to include snail migration with a constant migration rate, *ξ*, to and from an external population, ***N**^E^*, that is not affected by prawn interventions. For simplicity, the external population is conceptualized to originate from an identical transmission site to the one in which interventions are modeled, though in reality, multiple sites with heterogeneous transmission dynamics may contribute differentially to the intervention site.

**Figure 1:**
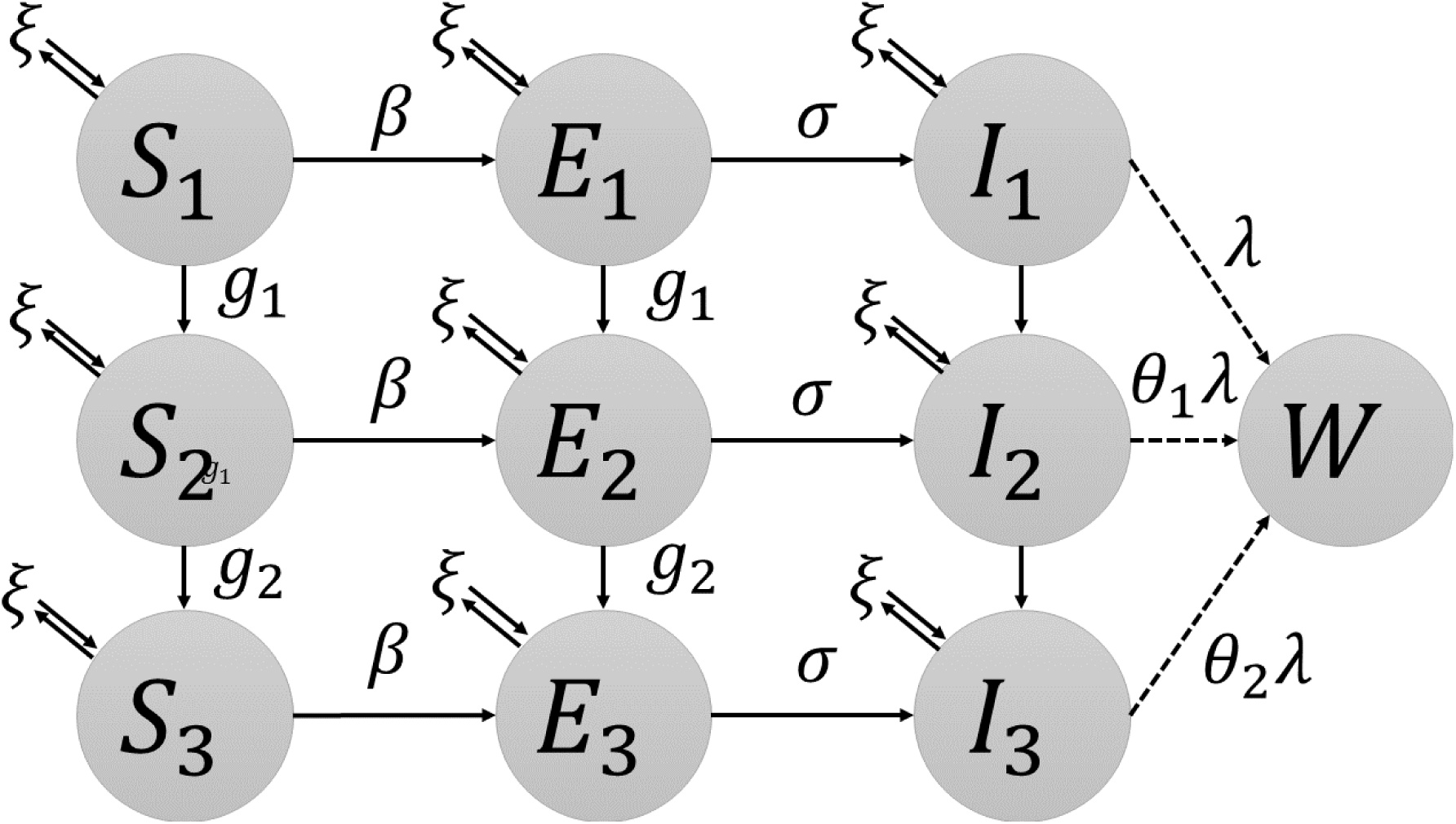
Epidemiologic model schematic with snail size and infection classes. Parameters governing transitions between classes and migration into and out of each class are shown.

New snails enter the population as susceptible juveniles, i.e. in infection class *i* = *S* and size class *j* = 1. Logistic snail population growth is modeled with *per*-*capita* recruitment, *f*, and carrying capacity, *K*. Neither small snails (*j* = 1) nor infectious snails (*i* = *I*) contribute to recruitment due to sexual immaturity and parasitic castration, respectively ^39^. Pre-patent snails’ contribution to recruitment is reduced by a fraction 0 < *z* < 1 ^40^. Snails of each size class are subject to both size-dependent natural mortality, *μ_j_*, and predation mortality, *ψ_j_*, a function of both prawn and snail body size and density described in the next section. Small and medium snails grow and transition to the next size class at the per capita rate *g*_1_ (from size class 1 to 2) and *g*_2_ (from class 2 to 3), respectively.

New snail infections occur at the per capita rate, *βM*, where *β* is the transmission rate and *M* = 0.5*ϕ*(*W*)*WHm* is an estimate of the overall number of *Schistosoma* miracidia (free living infective stages) produced by mated adult female worms. This estimate is the product of the size of definitive human host population, *H*, the mean parasite burden, *W*, (i.e., mean number of adult worms per person), the rate at which mated adult female worms shed eggs that hatch into infectious miracidia, *m*, and the function *ϕ*(*W*) representing the density-dependent mating probability of adult worms ^41^. The coefficient 0.5 accounts for the assumed 1:1 sex ratio of adult worms. For simplicity, we assume a constant human population and no density dependent fecundity of female worms. Following infection, pre-patent snails, *E*, transition to the patent class, *I*, at rate *σ*, with *σ*^−1^ being the mean time necessary for sporocyst development following snail infection with a miracidium.

The adult parasite population harbored by definitive human hosts is modeled as the mean parasite burden in the human population, *W*, assuming a negative binomial distribution with clumping parameter, *φ*^41,42^. Humans acquire adult worms as a result of contact with cercariae shed from infected snails. Worm acquisition occurs at rate *λ*, an aggregate parameter accounting for the *per capita* shedding rate of cercariae by infected snails, cercariae mortality, contact rate, and probability of infection, as described in previous work ^15^. The cercarial shedding rate of medium (*N*_*I*2_) and large (*N*_*I*3_) snails is assumed to exceed that of small (*N*_*I*1_) snails by a factor *θ*_1_ and *θ*_2_, respectively ^43^.

The full system of differential equations describing the epidemiologic model can be found in the supplementary information along with model parameters listed in Table S2.

### (c) Prawn predation model

As in previous work ^38^, the per-capita prawn predation rate on snails of each class, *ψ_ij_*, is modeled as a type III functional response, described by a generalization of Holling’s disk equation ^44^. This produces a sigmoid-shaped function, which increases and saturates at high prey densities and decreases to approach zero at low prey densities. Previous experiments by Sokolow et al ^13^ show that prawn predation of snails changes predictably as a function of the ratio of prawn biomass to snail body mass, *r_j_*. Using these experimental data, the attack rate, *α*, is estimated as a log-linear function of the biomass ratio: *α* = *α_m_* ∗ log(*r_j_*(*t*)). The handling time, *T_h_*, is estimated as a reciprocal function of the biomass ratio: *T_h_* = (*T_h_m__ r_j_*(*t*))^−1^ where *α_m_* and *T_h_m__* are coefficients estimated from ^13^ and *r_j_* the ratio between prawn body size and mean snail body size in each class. Laboratory experiments presented in 13 show that small prawns are unable to feed on large snails (*j* = 3), accordingly, *ψ* = 0 when *r_j_* < 3. In addition, the attack rate, *α*, derived by Sokolow et al. ^13^ in controlled laboratory conditions in 1*m*^−2^ tanks is penalized by a factor 0 < *ϵ* < 1 to account for the reduction in searching efficiency caused by the morphological complexity of foraging in wild settings.

The biomass ratio for each snail size class is estimated as:

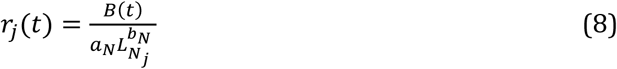

where *B*(*t*) is prawn body size derived with eq. (4), and the denominator represents snail mass in each class *j*, derived as a simple allometric function of snail shell diameter in each size class. The *per*-*capita* attack rate of prawns on snails of size *j* and infection class *i* is then estimated as:

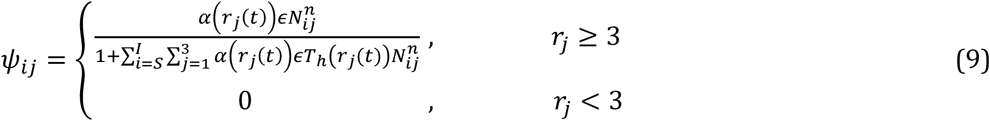

Prawn stocking at the considered densities is assumed to have no negative effects on water quality that may affect prawn survival, growth, predation of snails, or snail population dynamics, though ongoing field experiments in the lower Senegal River basin show that water quality may be negatively affected by nets installed to contain prawns introduced at transmission sites.

Parameters of the snail predation component of the combined model are listed in Table S3.

#### Model simulations

We consider a time horizon of *T_max_*= 10 years, for comparability to similar analyses investigating different schistosomiasis intervention strategies ^7,45^. Because prawn body size increases and levels off with time (eq. 1-3), but prawn abundance decreases in time (eq. 5), both stock biomass and cumulative profit (eq. 7) are unimodal functions of time for any given initial stocking density, *P*_0_. Additionally, harvesting prior to peak profit may afford the opportunity to increase *n^sp^* and therefore sacrifice short term profits to maximize long term profit. It is thus possible to use equations 1-7 to simulate prawn aquaculture dynamics and numerically find the stocking density, *P*_0_, and harvest time, *T*, that maximize cumulative profit. The surface of values (*P*_0_, *T*, *CP^sp^* (*P*_0_, *T*, *T_max_*)) is related to the eumetric curve used in fishery science to identify the stocking density that maximizes profit ^46^. As stocking costs increase linearly with stocking size, *P*_0_, while revenues increase less than linearly as a consequence of density-dependent growth and mortality, the surface is unimodal and its peak represents the maximum achievable cumulative profit, 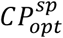 = max(*CP^sp^* (*P*_0_, *T*, *T_max_*)). Therefore, for each *Macrobrachium* species (*sp*) and a time period of *T_max_* = 10 years it is possible to identify the stocking density, 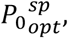 and harvest time, 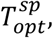 that maximize cumulative profit, 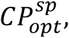 here collectively defined as *optimal management*. We identify optimal management for each prawn species using a grid search over initial stocking densities, *P*_0_, ranging between 0.5 – 7.5 *pm*^−2^ and potential harvest times, *T*, on each day between 1 – 730.

Given the optimal stocking density, 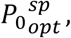 that maximizes cumulative profit and the corresponding optimal harvest time in an individual cycle, 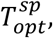 the epidemiologic model is run under the following scenarios: (1) 10 years of annual MDA with no prawn intervention; (2) and (3); 10 years of prawn stocking and harvesting under optimal aquaculture management for each species, as described above; and (4) and (5) 10 years of integrated annual MDA and prawn intervention under optimal management. In all scenarios, we simulate the system for an additional 10 years without intervention to explore the potential for infection rebound in the case the intervention program is ceased.

Stocking and harvesting were simulated at regular intervals of 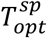 days, and were implemented as instantaneous events that reset the values of *P* and *L* to match the initial conditions at the beginning of each stocking cycle (i.e. 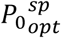 and *L*_0_). This assumes that all prawns—regardless of marketability—are harvested and replaced with 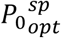 juveniles in a single day. Mass drug administration is implemented as an instantaneous 85% reduction in mean worm burden, *W*, in the same 75% of the human population, corresponding to assumptions of 85% drug efficacy and 25% systematic non-compliance ^45,47^.

To compare the effects of different interventions, we model disability associated with schistosomiasis using the disability adjusted life year (DALY) as in previous analyses ^7,48^. Disability weights measuring the disability associated with a condition for a single year of life—where 0 is perfect health and 1 is death—were distributed among individuals with heavy (> 50 eggs per 10mL urine, *H_hi_*) and light (0 < eggs per 10mL urine ≤ 50, *H_lo_*) *S. haematobium* burdens as defined by WHO guidelines. Total DALYs over the simulation period are then estimated as:

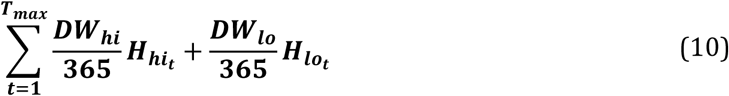

Where *DW_hi_* and *DW_lo_* are the yearly disability weights associated with heavy and light *S. haematobium* burdens, respectively, normalized to a daily estimate to match the dynamics of the epidemiological component of the model. Details on estimating the number of individuals in each burden class at each time step of the epidemiological model (*H_hi_t__* and *H_lo_t__*) are provided in the supplementary information.

The model was coded in R version 3.5.0 and simulated using the solver lsoda from the package deSolve ^49^. To address concerns regarding reproducibility, all model code and data are included as a supplementary file and are also made freely available online at https://github.com/cmhoove14/Prawn_fisheries_Public_health.

#### Sensitivity analyses

To quantify the influence of uncertain parameter inputs on key model outcomes, latin hypercube sampling is performed over parameter ranges in Table S1 to generate a set of 1,000 candidate sets. These parameter sets are used to derive estimates of uncertainty in the simulations described above. Furthermore, global sensitivity analysis using latin hypercube sampling and partial rank correlation coefficients (LHS-PRCC) ^50,51^ is performed with optimal cumulative profit 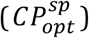 as the outcome in the prawn aquaculture model, equilibrium infected snail density (*N_I_*) and mean worm burden (*W*) absent prawn or MDA interventions as the outcomes in the epidemiologic model, and total DALYs accumulated through simulated 10-year combined MDA and prawn interventions as the outcome in the integrated model. Briefly, PRCC estimates the correlation between a model parameter and a model outcome by first rank-transforming parameters drawn from an LHS scheme and the corresponding model outputs, then removing the linear effects of all other model parameters on the parameter of interest and the outcome, and finally measuring the linear relationship between the rank-transformed parameter and outcome residuals ^50^. Monotonicity of the relationship between model outputs and all parameters was assessed via scatterplots (Figs S1-S4) prior to conducting LHS-PRCC sensitivity analyses.

Two additional sensitivity analyses are performed to identify profitability thresholds for the prawn aquaculture model. In the first, fixed costs, *c_f_*, in the range $0 – $1,000 are added to the per-cycle profit calculation (eqn 7; such that total cost for each cycle is –(*cP*_0_ + *c_f_*)) to represent additional costs that may be incurred with each aquaculture cycle such as labor, transport of prawns, and nursery costs leading up to stocking. In the second, the prawn mortality rate, *μ_P_*, is multiplied up to 10 times the estimated natural rate to account for additional sources of prawn loss that may be incurred due to predation, escape, or other sources of mortality, such as disease.

## Results

### Aquaculture model

With parameters fixed at the values shown in Table 1, stocking *M. rosenbergii* at *P*_0_ = 2.6 *Pm*^−2^ and harvesting at 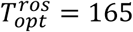 *days* maximizes cumulative ten-year profit while stocking at *P*_0_ = 2.4 *Pm*^−2^ and harvesting at 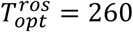 *days* maximizes cumulative profit for *M. vollenhovenii*. These stocking densities and harvesting times were used to simulate aquaculture cycles for each species. Figure 2 shows the dynamics of each species run continuously through two years with vertical dashed lines indicating the optimal time of harvest, 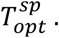 Prawns grow in length, *L*, and weight over time while population size (measured as density, *Pm*^−2^) decreases with time as a result of density dependent death from crowding and natural, size-dependent mortality (Fig 2). These competing effects lead to a humped function of total harvestable biomass, *Ω*, over time with the peak occurring well before prawns grow to their full size. Ten-year cumulative profits also have a single peak, which is determined both by the profit per cycle and the number of cycles possible within the 10 year time frame. Cumulative profits are maximized by harvesting well before the peak in harvestable biomass occurs, indicating more, smaller harvests maximize profit over time.

**Figure 2:**
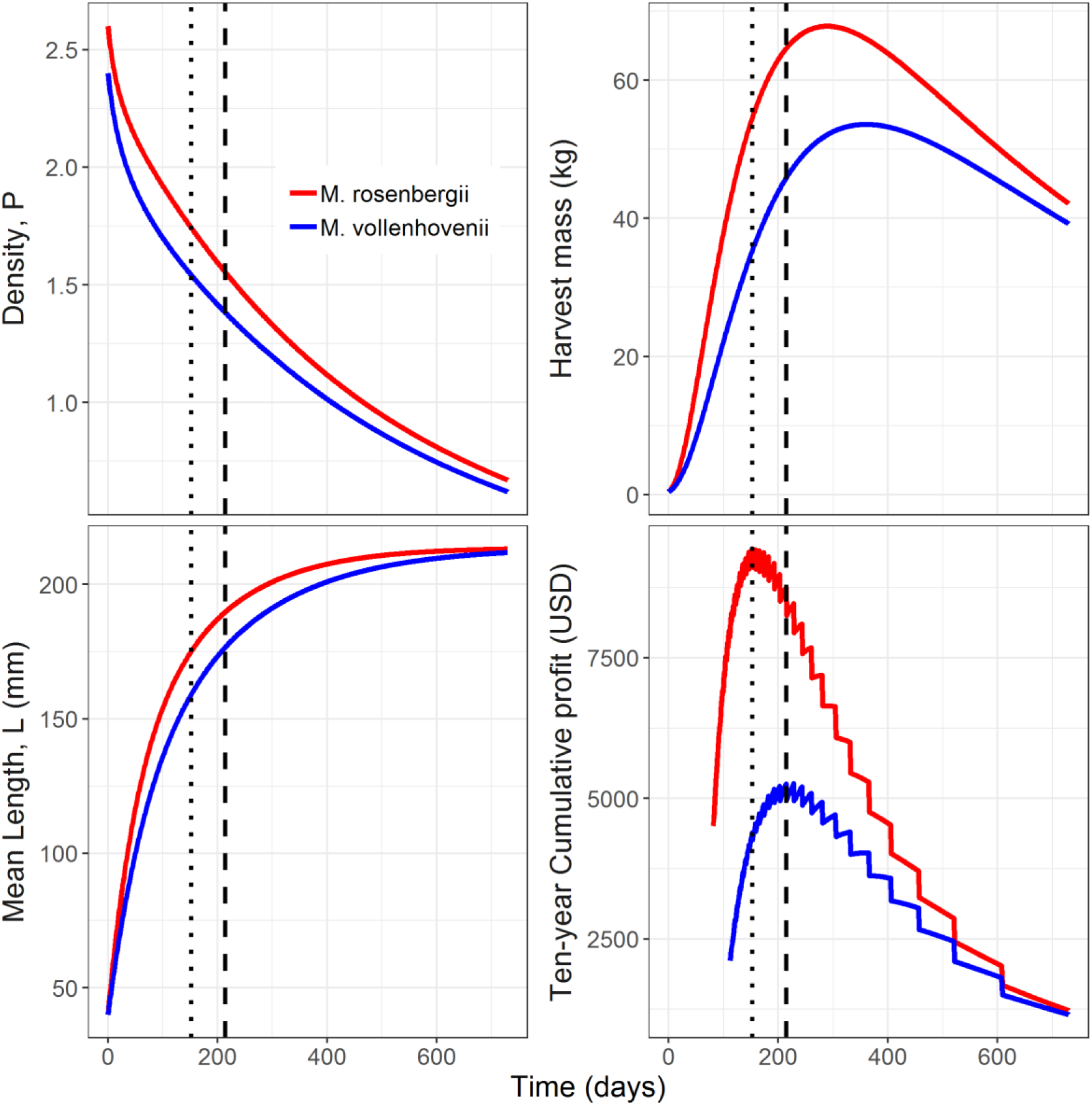
Prawn aquaculture model dynamics. Two-year aquaculture cycles for, *M. rosenbergii* (red lines) and *M. vollenhovenii* (blue lines) under optimal management showing how prawns grow in length over time (bottom-left), but decrease in density (top-left). This leads to a single peak in harvest mass (top-right), but harvesting actually occurs prior to the peak in order to maximize ten-year cumulative profits (bottom-right) by sacrificing profit-per-cycle for completing more aquaculture cycles. Vertical dashed lines indicate time at which harvest would occur for each species (small dashes – *M. rosenbergii*, large dashes – *M. vollenhovenii*). Here, following (19), we set the cost *c* for a juvenile prawn to $0.10 with *L*_0_ = 40*mm* (~0.35g per juvenile prawns) and selling price p=$12/kg. Other parameters set as in Table 1.

**Table 1:** Optimal stocking and harvesting parameters for each prawn species reported as median (interquartile range).

| **<u>Parameter</u>** | **<u>Definition</u>** | **<u>*M. vollehovenii*</u>** | **<u>*M. rosenbergii*</u>** |
| --- | --- | --- | --- |
| $P_{0opt}^{sp}$ | Optimal stocking density | $2.4 Pm^{-2}$ | $2.6 Pm^{-2}$ |
| $L_0$ | Stocking size of juveniles | 40mm | 40mm |
| $T_{opt}^{sp}$ | Optimal harvest time | 260 days<br>(228 – 331) | 165 days<br>(146 – 192) |
| $L(T_{opt}^{sp})$ | Mean length at harvest | 167mm<br>(161 – 175) | 167mm<br>(161 – 174) |
| $P(T_{opt}^{sp})$ | Prawns harvested | 1056<br>(850 – 1236) | 1559<br>(1434 – 1662) |
| $Y(T_{opt}^{sp}, P_{0opt}^{sp})$ | Commercial yield per harvest | 28 kg<br>(21 – 36) | 41 kg<br>(34 – 49) |
| $n^{sp}$ | Number of cycles in 10 years | 14<br>(11 – 16) | 22<br>(19 – 25) |
| $CP_{opt}^{sp}$ | Cumulative profits over 10 years | \$1891<br>(\$856 – \$3486) | \$5403<br>(\$3380 – \$8075) |

The surface of values (*P*_0_, *T*, *CP^sp^*(*P*_0_, *T*, *T_max_*)) for each species is shown in Figure 3. As expected, profits associated with aquaculture of the faster growing *M. rosenbergii* are higher. Considering parametric uncertainty, the peak estimate of median cumulative profit for *M. rosenbergii* occurs at *P*_0_ = 2.9 *Pm*^−2^ and 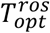 = 173 *days* (IQR: 146 – 192), producing 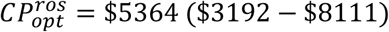 per 1,000 *m*^2^ enclosure. The same estimates for *M. vollenhovenii* are *P*_0_ = 2.5 *Pm*^−2^, 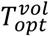 = 260 *days* (228 – 331), and 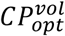 = $1738 ($704 – $3394). Additional outputs from the aquaculture model that describe stock structure and aquaculture performances at optimal management are reported in Table 1.

**Figure 3:**
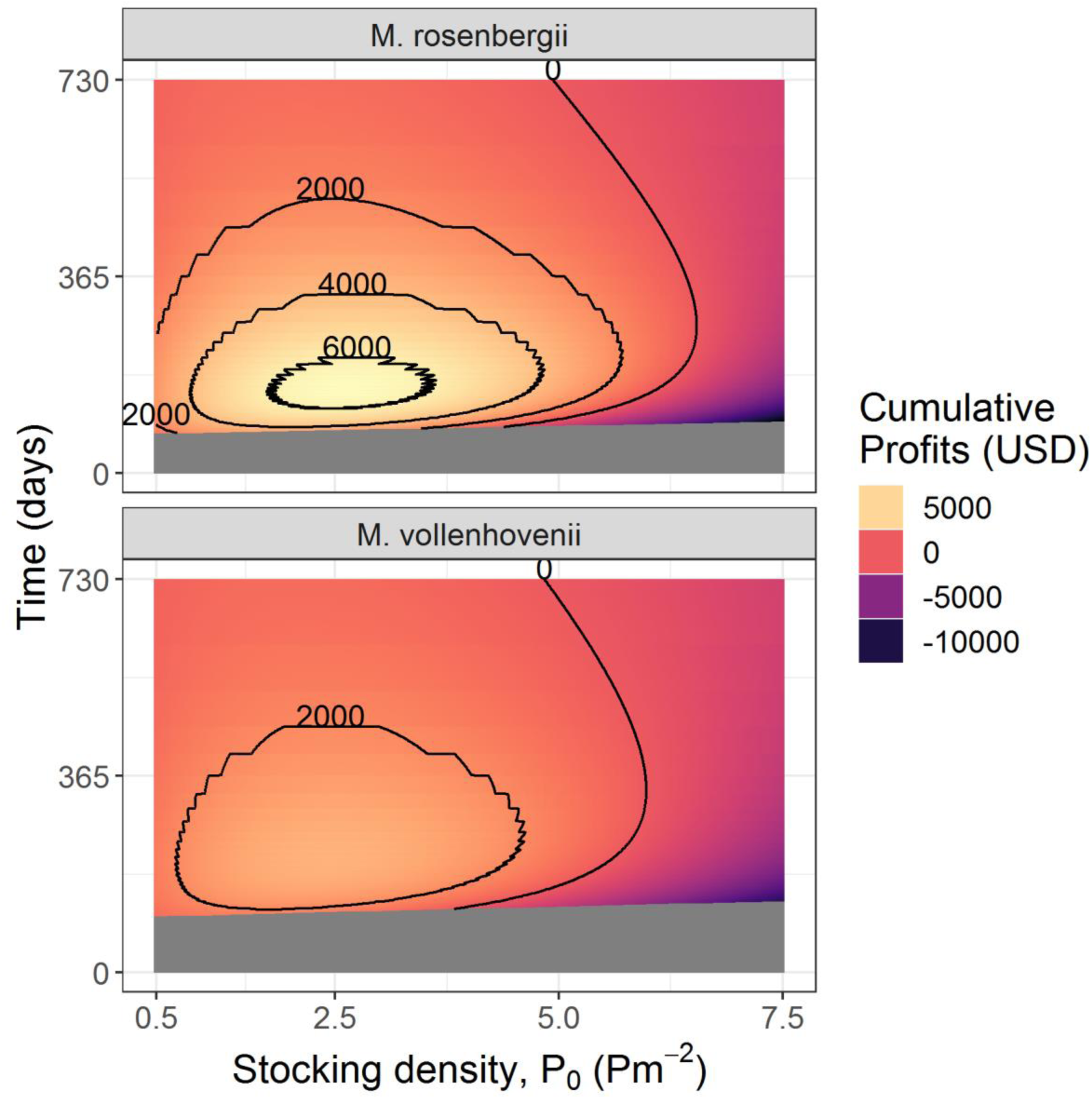
Grid search to identify optimal management decisions for each prawn species. Cumulative profits (*CP^sp^*) generated by the aquaculture model for each species across a range of potential stocking densities (*P*_0_) and harvest times (*T*) are shown. Grey regions indicate regions where harvesting is not feasible due to prawns not having reached sufficient marketable size (30*g*). Contours indicate regions of *CP^sp^* corresponding to the labeled value in 2018 USD.

### Integrated aquaculture epidemiologic model

Simulation of the integrated model reveals comparable performance of prawn intervention strategies to MDA strategies. As in previous analyses ^7,15^, annual mass drug administration alone produces a pattern of gradually decreasing worm burden over time characterized by repeated rebound in infection following MDA (Fig 4A, purple line). The prawn-only intervention causes mean worm burden to gradually decline towards 0, eventually reducing worm burden to comparable levels by year 10 (Fig 4A, blue line). Finally, the combined MDA and prawn intervention leads to a rapid decline in mean worm burden that nearly reaches 0 by year 10 (Fig 4A, gold line).

**Figure 4:**
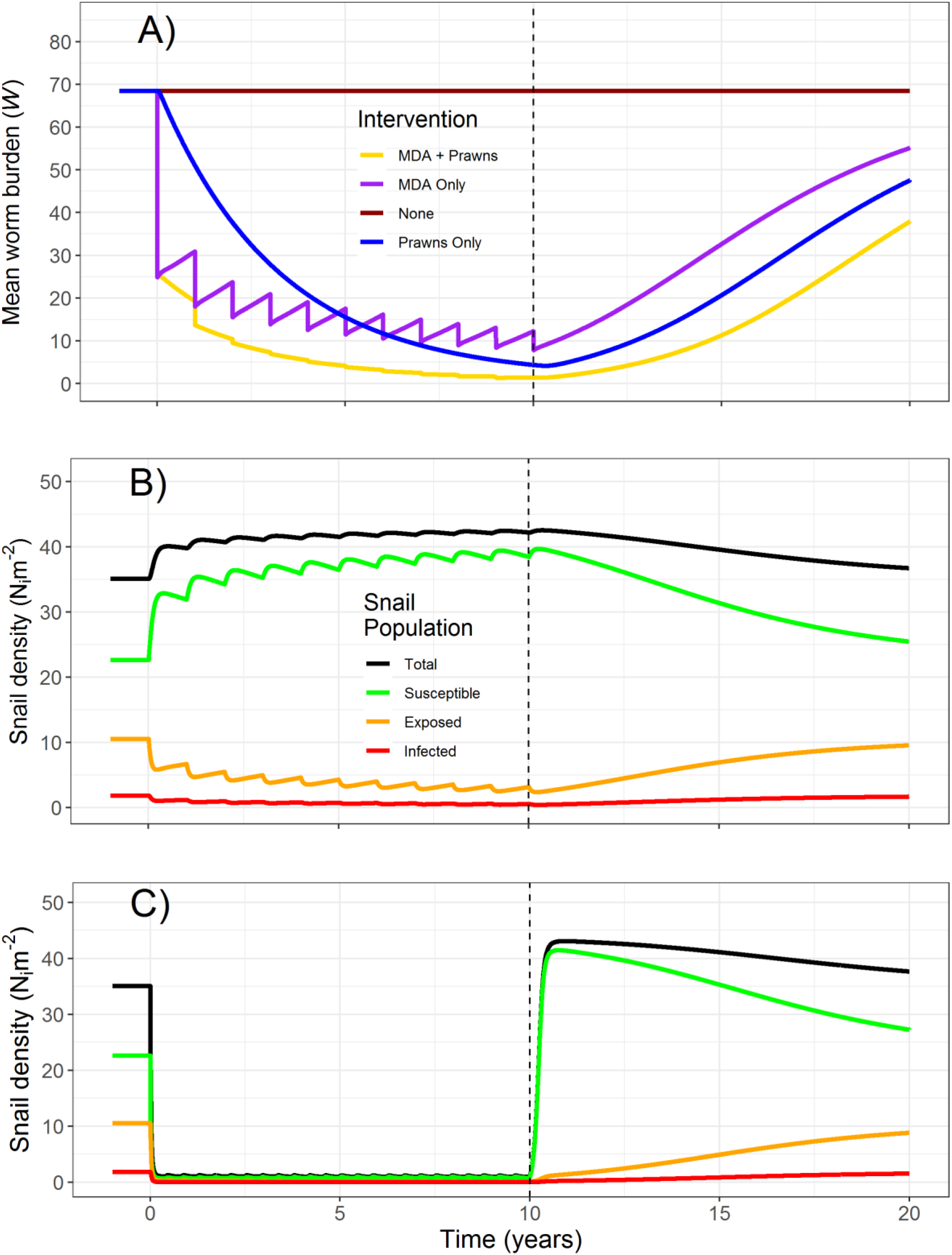
Outputs of the integrated model under different intervention scenarios implemented over ten years followed by ten years of no intervention. Worm burden trajectories under no intervention (red), annual MDA only (purple), prawn stocking of *M. rosenbergii* under optimal management (blue), and both annual MDA and prawn stocking (gold) (A); snail infection dynamics under MDA only intervention (B); and snail infection dynamics under prawn stocking interventions (C). Outputs from *M. volenhovenii* interventions not shown as they approximately mirror those of *M. rosenbergii*.

These patterns can mostly be explained by the underlying snail infection dynamics. Under MDA intervention, the snail population persists through repeated rounds of MDA and even increases due to the reduced influence of infection (Fig 4B). Most importantly, a population of infected snails is able to sustain transmission-albeit at lower levels-even as adult worms are removed from the treated human population by MDA (Fig 4B). Interventions in which prawns are introduced at profit-optimal densities produce rapid declines in the snail population that heavily reduce this environmental reservoir of transmission (Fig 4C). Extirpation of the entire snail population is prevented due to the assumption of a Holling’s type III functional response and immigration from an unaffected reservoir population (see Fig S3 for snail infection dynamics without immigration and under a Holling’s type II functional response), but transmission is effectively halted due to near elimination of the infected snail population (Fig 4C). This heavy reduction in transmission coupled with the benefits of MDA translates to near elimination of the parasite by year 10. Regardless of the intervention, ceasing efforts to control transmission after 10 years results in rapid returns to pre-intervention snail populations and community-level mean worm burdens (Fig 4A-C).

Comparison of total DALYs lost over ten year simulation periods under each intervention shows comparable performance of the prawn only intervention to MDA and ubstantial additional DALYs averted when combining MDA with prawn intervention. Without intervention a median 324 (IQR: 119 – 502) DALYs are lost to *S. haematobium* infection. Annual MDA and profit-optimal stocking of prawns perform comparably with 160 (IQR: 54 – 285) total DALYs lost with annual MDA and 184 (IQR: 70 – 294) total DALYs lost with profit-optimal prawn stocking; representing 51% and 43% DALYs averted, respectively. Integrated interventions utilizing both MDA and prawn stocking reduce median DALYs lost to 83 (IQR: 30 – 137), representing a 74% reduction in *S. haematobium* related disability.

### Sensitivity Analyses

Substantial sensitivity analyses reveal that profitability of the prawn aquaculture system and reductions in schistosomiasis burden associated with prawn introductions are robust to model assumptions and parameters (Figs S1 and S2). The addition of fixed costs associated with each aquaculture cycle and additional sources of mortality reveal aquaculture of *M. rosenbergii* remains profitable with fixed costs of up to $550 per cycle (Fig S1) and mortality rates up to 5.2 times higher than the baseline estimate (Fig S2). Aquaculture of *M. vollenhovenii* remains profitable with fixed costs of up to $400 per cycle (Fig S1) and mortality rates up to 3.6 times higher than the baseline estimate (Fig S2). Global sensitivity analysis using LHS-PRCC shows that the parameters of the VBGE that govern prawn growth dynamics (*L*_∞_, *κ*) and parameters governing estimates of discounted profits (*δ*, *p*) are most correlated with 10-year cumulative profits (Fig 5B)

**Figure 5:**
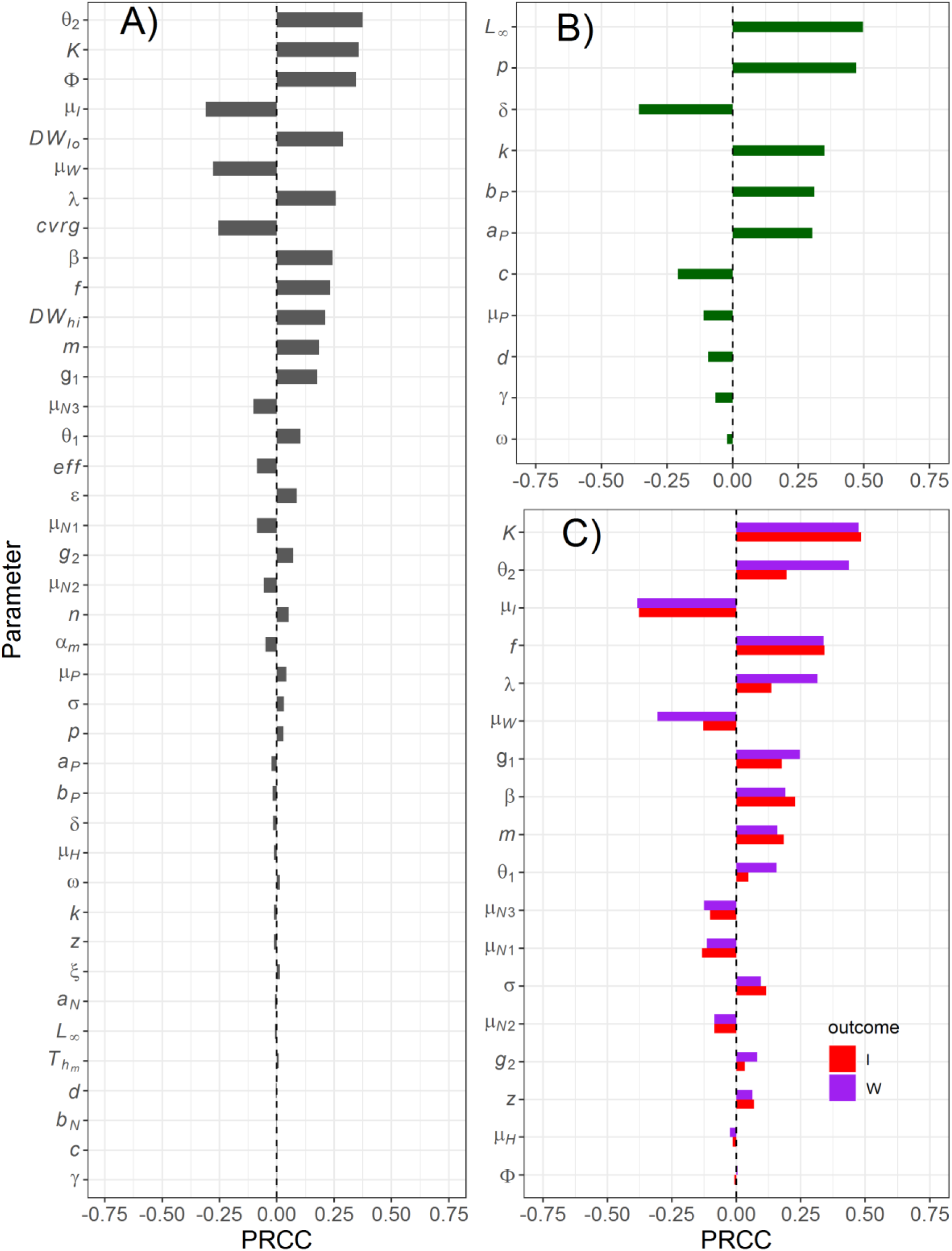
Sensitivity of key model outcomes to model parameters assessed with LHS-PRCC. Sensitivity to all parameters of DALYs lost following 10 years of combined MDA and prawn intervention (A), cumulative 10-year profit sensitivity to prawn aquaculture parameters (B), and sensitivity of mean worm burden and equilibrium infected snail density to epidemiologic parameters (C). Parameter definitions and baseline values can be found in supplementary tables 1-3.

Exploration of snail infection dynamics under prawn interventions also reveals that the Holling’s type III functional response and the inclusion of snail migration represent conservative structural model assumptions. Figure S3 shows that snail extirpation is only possible when both of these assumptions are relaxed. Parameters governing snail predation by prawns (*ϵ*, *n*) are also shown to have moderate influence on estimates of 10-year cumulative DALYs lost (Fig 5A).

Key outcomes from the epidemiologic model including equilibrium estimates of infected snail density (*I*) and mean worm burden (*W*) and of DALYs lost over 10-year intervention periods is mostly influenced by epidemiologic parameters, rather than parameters governing prawn population or predation dynamics (Fig 5A & 5C). We also compared parameter sets in which the MDA only intervention was superior to the prawn only intervention in terms of DALYs averted. We find that the MDA only intervention leads to more DALYs averted when the coverage and efficacy are high, while the prawn-only intervention performs better when the snail population carrying capacity (*K*) is higher (Fig S4).

## Discussion

Small-scale, extensive prawn aquaculture such as that considered here offers a profitable and sustainable method to improve food production, reduce schistosomiasis burdens, and increase revenues for small-scale subsistence farmers, especially when paired with ongoing efforts such as rice cultivation ^25^. In areas of high food insecurity, malnutrition, and endemic schistosomiasis—conditions which reinforce each other and often result in “poverty traps” ^52,53^—an integrated system of prawn aquaculture may be a solution for co-benefits of disease control and sustainable development.

Our results are consistent with an increasing body of evidence that consumption of snails by predator species such as *Macrobrachium* prawns can be an effective method for combating schistosomiasis transmission to people ^14,15,54^. Specifically, by targeting the environmental reservoir of schistosome parasites, prawns can reduce reinfection rates that plague MDA campaigns in high transmission settings. By deploying prawn aquaculture with MDA, effective control—in which the schistosome parasite is suppressed in both human hosts and intermediate host snails—can be achieved.

We also show that such prawn interventions combined with established extensive aquaculture methods can be profitable if carefully managed. Drawing on economic studies of existing aquaculture practices ^19,33^, we develop—to our knowledge—the first dynamic model of prawn aquaculture to investigate optimal management practices. The model suitably simulates prawn stocking in a large water contact site or rice paddy, and the length of optimal cycles coincides with typical rice harvesting timelines ^25^. We find that this system produces short- and long-term profits, implying the potential for a sustainable, community-driven intervention, given the right capital investment and incentive programs.

Under optimal management, extensive aquaculture of of either *M. rosenbergii* or *M. vollenhovenii* leads to both profits and reductions of the snail population. The optimal stocking densities of each species are above the potential threshold of approximately 2 *Pm*^−2^ necessary for local snail extirpation as identified in previous work ^15^, though our conservative inclusion of snail migration and a Holling’s type III functional response restricts such an outcome in this analysis.

Achieving these stocking densities may be challenging in sub-Saharan Africa where >90% of the global burden of schistosomiasis is found ^23^ and where the proposed intervention is likely to provide the most benefit. More than 50 years of domestication of *M. rosenbergii* has established protocols for ideal rearing and management of aquaculture efforts with this species ^18,19^, whereas similar protocols for the African native *M. vollenhovenii* are still under development. Profits appear to be highly sensitive to parameters that regulate prawn growth, meaning continued research and development into *M. vollenhovenii* aquaculture may eventually provide comparable profits to *M. rosenbergii*. However, in the short term, all male stocking of *M. rosenbergii* may be a superior strategy.

This strategy of stocking the non-native *M. rosenbergii* in coastal regions where it is non-native should be approached with caution to avoid the establishment of an alien population with potentially unintended ecological consequences on local biodiversity. Establishment of non-GMO aquaculture biotechnologies for either all-male ^55-57^ or all-female ^58^ populations suggest possible strategies to prevent invasions. Moreover, recent laboratory experiments (Savaya-Alkalay et al. submitted) have ruled out the possibility of cross-fertilization between mature female *M. vollenhovenii* and male *M. rosenbergii*, demonstrating that male-only progeny of *M. rosenbergii* may be safely used for extensive aquaculture and schistosomiasis control in western Africa where *M. rosenbergii* is non-native.

Extensive sensitivity analyses reveal that key model outcomes including profit generated by the aquaculture portion of the model and DALYs lost from the integrated model are robust to key parameters that govern profit estimation and the influence of prawn interventions on DALYs lost. The prawn aquaculture model is most sensitive to parameters governing somatic growth, which have been estimated from available literature, and supported by expert opinion ^28,32^. This finding also suggests that if stock improvement or improvements in husbandry practices can increase average prawn condition or size, i.e. by supplementing feeds, further increases in profit may be possible. Selective harvesting methods that only remove market-size prawns, leaving the remaining stock to continue to grow, may also improve aquaculture performance.

Optimal aquaculture practices are also sensitive to the market price of adult prawns and the stocking cost of juvenile prawns, implying that optimal management may be influenced by fluctuations in market factors that may influence price and cost of harvested and stocked prawns. Profit generated from selling harvested prawns is based on reasonable assumptions of prices in premium markets, though the actual selling price in subsistence economies might be lower and contingent on market access and local demand. However, the fixed cost sensitivity analysis demonstrates profitability is possible even with substantial additional costs associated with prawn stocking, and profitability persists across a wide range of stocking densities for each species. These results together assuage concerns that such vagaries of the market would impair the success of the proposed system and suggest that prawn aquaculture should be economically viable even under non-optimal management densities.

Additional sources of mortality could result from declines in water quality caused by prawn stocking, disease outbreaks, or predation by waterbirds, fish, amphibians or reptiles. We demonstrate that extensive aquaculture is still profitable with mortality rates as high as five times greater than natural mortality. Barring catastrophic events, this demonstrates that the proposed system should be resilient to such perturbations. Therefore, further field work is required to assess prawn life expectancy, escapement rate, growth performances, changes in water quality, and potential changes in human behavior which might affect either aquaculture performances and/or snail abundance, transmission rates, and epidemiologic outcomes.

Prawn predation of the snail population as modeled here is also based on a number of assumptions including the Holling’s type III functional response and the prawn attack rate penalty. We consider the type III functional response a conservative estimate of the relationship between prawn predation and snail density as it does not allow for the potential extirpation of the snail population. We also test a wide range of attack rate penalties as we identified no prior estimates of prawn predation efficiency in non-laboratory settings, and find that effective control is feasible even when increasing this parameter by an order of magnitude (implying a substantially reduced attacked rate).

Regarding schistosomiasis transmission dynamics, recent findings suggest non-linearities in the human-to-snail force of infection may decrease the efficacy of MDA and lead to faster post-MDA rebounds of schistosomiasis ^59^. While this may alter the effects of MDA in our simulations, we believe it strengthens the argument for strategies that explicitly target the intermediate host snail population, such as the prawn intervention proposed here. Another recent finding suggests that snail control can actually lead to increased human risk of *Schistosoma* infection if the snail population is limited by resource competition prior to “weak” control efforts ^60^. In this scenario, remaining snails with higher per-capita resource availability may produce more cercariae. Given the large magnitude of snail reductions at the proposed prawn densities modeled here—even given the conservative Holling’s III functional response—we believe this effect is unlikely in our scenarios. Finally, our model lacks seasonality, which would likely affect both schistosomiasis transmission and prawn growth as e.g. temperature and rainfall fluctuate through the year, especially in sub-tropical and near temperate regions where schistosomiasis is still endemic, such as in northern Africa ^61,62^.

This bioeconomic analysis shows that an integrated intervention strategy that utilizes both MDA and a profitable prawn aquaculture system can successfully control schistosomiasis while generating profit. Since the intervention is driven by a profitable business model, it may be sustainable purely through market incentives, and thereby reduce the need for external support from donors or public health agencies. Subsidies are only likely to be necessary in the event that *M. rosenbergii* aquaculture is not suitable for the region and obtaining large quantities of *M. vollenhovenii* juveniles is infeasible or expensive. Research and development for this system is indeed ongoing in Senegal, which will aid future analyses of the effectiveness and feasibility of this promising integrated strategy.

## Data Availability

All data and code used to conduct this analysis are provided as a supplementary file and are freely available at https://github.com/cmhoove14/Prawn_fisheries_Public_health

## Acknowledgements

CMH, JVR, GADL, IJJ, AJL, SHS, and JRR were supported by the National Institutes of Health grant no. R01TW010286 (to JRR and JVR). CMH and JVR were additionally supported in part by the National Science Foundation Water, Sustainability and Climate grants 1360330 and 1646708 (to JVR), by National Institutes of Health grant no. R01AI125842 (to JVR), and by the University of California Multicampus Research Programs and Initiatives award MRP-17-446315 (to JVR). GADL, IJJ, AJL and SHS were additionally funded by a grant from the Bill and Melinda Gates Foundation (OPP1114050) and by a GDP SEED grant from the Freeman Spogli Institute at Stanford University. GADL, IJJ, AJL, SHS, and JNS were also supported by NSF CNH grant #1414102. GADL, SHS, CMH, JVR, JNS, RC, LM and MG were supported also by NIMBioS through the working group on the Optimal Control of Environmentally Transmitted Disease. JPS and AR acknowledge funds provided by the Swiss National Science Foundation, via the project “Optimal control of intervention strategies for waterborne disease epidemics” (200021-172578/1). CLW was supported by the Michigan Society of Fellows at the University of Michigan and by a Sloan Research Fellowship from the Alfred P. Sloan Foundation. RC and LM were also supported by Politecnico di Milano through the Polisocial Award programme (project MASTR-SLS).

## Author information

### Contributions

GADL and SHS conceived the problem and designed the general modelling framework. CMH, SHS, JK, JVR, and GADL developed the analysis. CMH and JK wrote the simulations scripts. GR collected field data to parameterize the epidemiologic model. SHS provided experimental data to parameterize the predation component of the model. JNS provided guidance on profit estimation of the prawn aquaculture model. AS-A, SC, and AS provided guidance on dynamics of the aquaculture model. CMH, JK, JNS, JVR, and GADL drafted the manuscript and all authors contributed to its editing.

## Supplementary Information

**Table S1:** Parameters of the prawn aquaculture model and ranges used in the global sensitivity analysis

| Parameter | Value & Units | Description | Source | Range |
| --- | --- | --- | --- | --- |
| $L_{\infty}$ | 214 mm | Maximum length used in von Bertalanffy growth equation | 32 | [184, 234] |
| $\kappa$ | $0.0034 \frac{\text{mm}}{\text{day}}$ | Growth rate of *Macrobrachium vollenhovenii* | 32,63 | [0.0078, 0.012] |
| | $0.0104 \frac{\text{mm}}{\text{day}}$ | Growth rate of *Macrobrachium rosenbergii* | 29 | |
| $\gamma$ | $3.5 \times 10^{-6} g^{-1}$ | Density dependent growth reduction | 26 | $[3.5 \times 10^{-6}, 6.5 \times 10^{-6}]$ |
| $a_p$ | 0.00244 | Allometric parameter for *Macrobrachium* length-weight conversion | 28 | [0.066, 0.081] |
| $b_p$ | 3.55 | Allometric parameter for *Macrobrachium* length-weight conversion | | [3.46, 3.64] |
| $\mu_p^{\dagger}$ | $0.0061 \frac{P}{\text{day}}$ | Daily prawn mortality rate | 32,63 | [0.0054, 0.012] |
| $d$ | -0.382 | Size-dependent mortality scaling coefficient | 31 | [-0.461, -0.289] |
| $\omega$ | $5.5 \times 10^{-9} g^{-1}$ | Density dependent mortality factor | 26 | $[7.5 \times 10^{-9}, 3.5 \times 10^{-9}]$ |
| $p$ | $\frac{\$12}{kg}$ | Price per kg of harvested market-size prawns | 33,64,65 | [11, 22] |
| $\delta$ | $\frac{1.99 \times 10^{-4}}{\text{day}}$ | Discount rate | (7% annual) | [3%, 20%] |
| $c^{\dagger}$ | $\frac{\$0.10}{P}$ | Cost of a stocked juvenile prawn | 33 | [0.045, 0.12] |

<sup>†</sup> See additional sensitivity analyses below

## Epidemiologic model equations

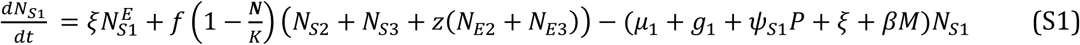

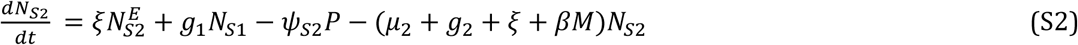

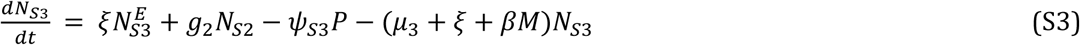

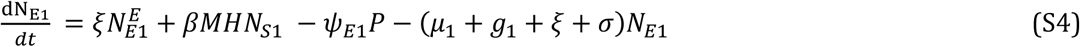

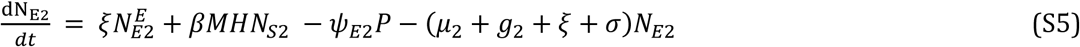

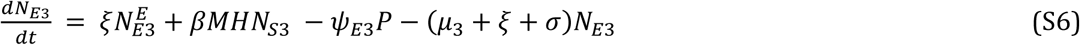

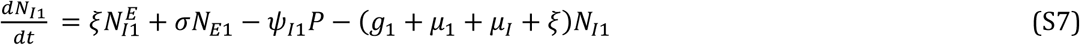

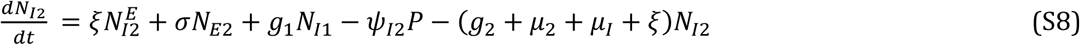

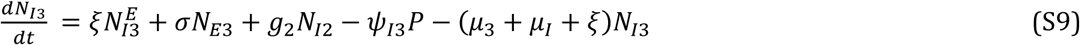

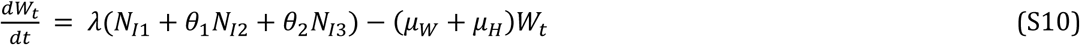

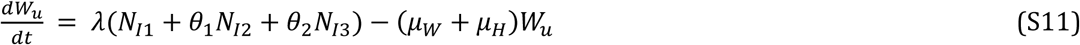

## MDA Implementation and DALYs estimation

The model tracks treated and untreated segments of the population by dividing the worm burden based on MDA coverage, such that total worm burden is a weighted average of each population: *W* = *cvrgW_t_* + (1 – *cvrg*)*W_u_*. When MDA is implemented, mean worm burden in the treated compartment is reduced by *W_t_* (1 – *eff*). We assume the dispersion parameter of the negative binomial distribution in each population is constant over the simulation period. To estimate the total number of individuals with heavy and light egg burdens, *H_hi_* and *H_lo_*, respectively, we first sample *n* = *H* ∗ *cvrg* draws from a negative binomial distribution with mean *W_t_* and dispersion, *φ*, to represent the distribution of adult worms in the treated segment of the human population. Similarly, we sample *n* = *H* ∗ (1 – *cvrg*) draws from a negative binomial distribution with mean *W_u_* and dispersion, *φ*, to represent the distribution of adult worms in the untreated segment of the human population. We then convert these individual level adult worm counts, denoted *W_h_*, to estimates of egg burden, denoted *B_h_*, as: *B_h_* = 0.5*W_h_ϕ*(*W_h_*)*ε* where 0.5*W_h_ϕ*(*W_h_*) provides an estimate of the number of mated female (i.e. egg producing) worms and *ε* is an estimate of *S. haematobium* eggs per 10mL urine per mated adult female worm from [CITE]. With the full distribution of *B_h_*, we can then estimate the number of individuals with heavy (> 50 eggs per 10mL urine, *H_hi_*) and light (0 <eggs per 10mL urine ≤ 50, *H_lo_*) *S. haematobium* burdens as defined by WHO guidelines and subsequently estimate DALYs as in equation 10.

**Table S2:** Parameters of the schistosomiasis epidemiologic model

| Parameter | Value & Units | Description | Source | Range |
| --- | --- | --- | --- | --- |
| $f$ | $0.26 N_{S1} day^{-1}$ | Per-capita recruitment rate of reproductive snails | 15 | [0.06, 0.36] |
| $K$ | $50 \frac{N}{m^2}$ | Snail population carrying capacity | 66 | [25, 75] |
| $z$ | 0.5 | Proportion of pre-patent snails that reproduce | 40 | [0.25, 1] |
| $\mu_1$ | $\frac{1}{50} day^{-1}$ | Mortality rate of snails in size class 1 | 67,68 | [25, 100] |
| $\mu_2$ | $\frac{1}{75} day^{-1}$ | Mortality rate of snails in size class 2 | | [50, 125] |
| $\mu_3$ | $\frac{1}{100} day^{-1}$ | Mortality rate of snails in size class 3 | | [75, 150] |
| $\mu_I$ | $\frac{1}{10} day^{-1}$ | Mortality rate of snails in infection class $I$ | 40 | [7, 20] |
| $\xi$ | $\frac{1}{5} year^{-1}$ | Migration rate of snails | 69 | [0, 1] |
| $g_1$ | $\frac{1}{37} day^{-1}$ | Growth rate of snails in size class 1 | 70 | [20, 60] |
| $g_2$ | $\frac{1}{62} day^{-1}$ | Growth rate of snails in size class 2 | | [40, 100] |
| $\sigma$ | $\frac{1}{50} day^{-1}$ | Pre-patent period; transition rate of snails in infection class $E$ to infection class $I$ | 71 | [30, 70] |
| $m^\dagger$ | 0.8 | Infectious miracidia produced per mated adult female worm | 15 | [0.5, 1.5] |
| $\beta^\dagger$ | $4 \times 10^{-7}$ | Pre-patent snails produced per infectious miracidia | | $[2.0 \times 10^{-7}, 8 \times 10^{-7}]$ |
| $\lambda^\dagger$ | $7.5 \times 10^{-6}$ | Adult worms produced per infectious cercaria shed by infected snails | | $[5.0 \times 10^{-6}, 1.0 \times 10^{-5}]$ |
| $\theta_1$ | 1.31 | Increased relative production of infectious cercariae by snails in class $N_{I2}$ | 43 | [1, 5] |
| $\theta_2$ | 7.88 | Increased relative production of infectious cercariae by snails in class $N_{I3}$ | | [2, 10] |
| $\varphi$ | 0.08 | Clumping parameter of the negative binomial distribution | 38 | [0.01, 0.3] |
| $\mu_w$ | $\frac{1}{3.3} year^{-1}$ | Mortality rate of adult worms | 15 | [2, 4] |
| $\mu_H$ | $\frac{1}{60} year^{-1}$ | Mortality rate of humans harboring adult worms | | [50, 70] |
| $DW_{hi}$ | 0.05 | Disability weight associated with heavy *S. haematobium* egg burden | 7,72 | [0.03, 0.15] |
| $DW_{lo}$ | 0.014 | Disability weight associated with heavy *S. haematobium* egg burden | 7,72 | [0.003, 0.03] |
| $eff$ | 0.85 | MDA efficacy | | [0.75, 0.95] |
| $cvr g$ | 0.75 | MDA coverage | | [0.5, 0.95] |
| $\mathcal{E}^\dagger$ | 3.6 | Eggs produced per 10mL urine per mated adult female worm | 73 | [2, 5] |

† Units provided in description

**Table S3:** Parameters regulating snail predation of snails, linking the aquaculture model to the epidemiologic model

| Parameter | Value & Units | Description | Source | Range |
| --- | --- | --- | --- | --- |
| $a_N$ | 0.19 | Allometric parameter for snail shell width-weight conversion | 13 | [0.1, 0.3] |
| $b_N$ | 2.54 | Allometric parameter for snail shell width-weight conversion | | [2, 3] |
| $\alpha_m$ | 0.91 | Parameter to estimate prawn attack rate from prawn-snail biomass ratio | | [0.5, 1.5] |
| $T_{hm}$ | 0.39 | Parameter to estimate prawn handling time from prawn-snail biomass ratio | | [0.2, 0.5] |
| $n$ | 2 | Exponent of Holling's type III functional response | 38 | [1.1, 4] |
| $\epsilon$ | 10% | Prawn predation attack rate penalty associated with searching for prey in wildlife rather than laboratory conditions | Estimated as 10% of laboratory-determined rates with wide range investigated in sensitivity analysis | [1, 100] |

**Figure S1:**
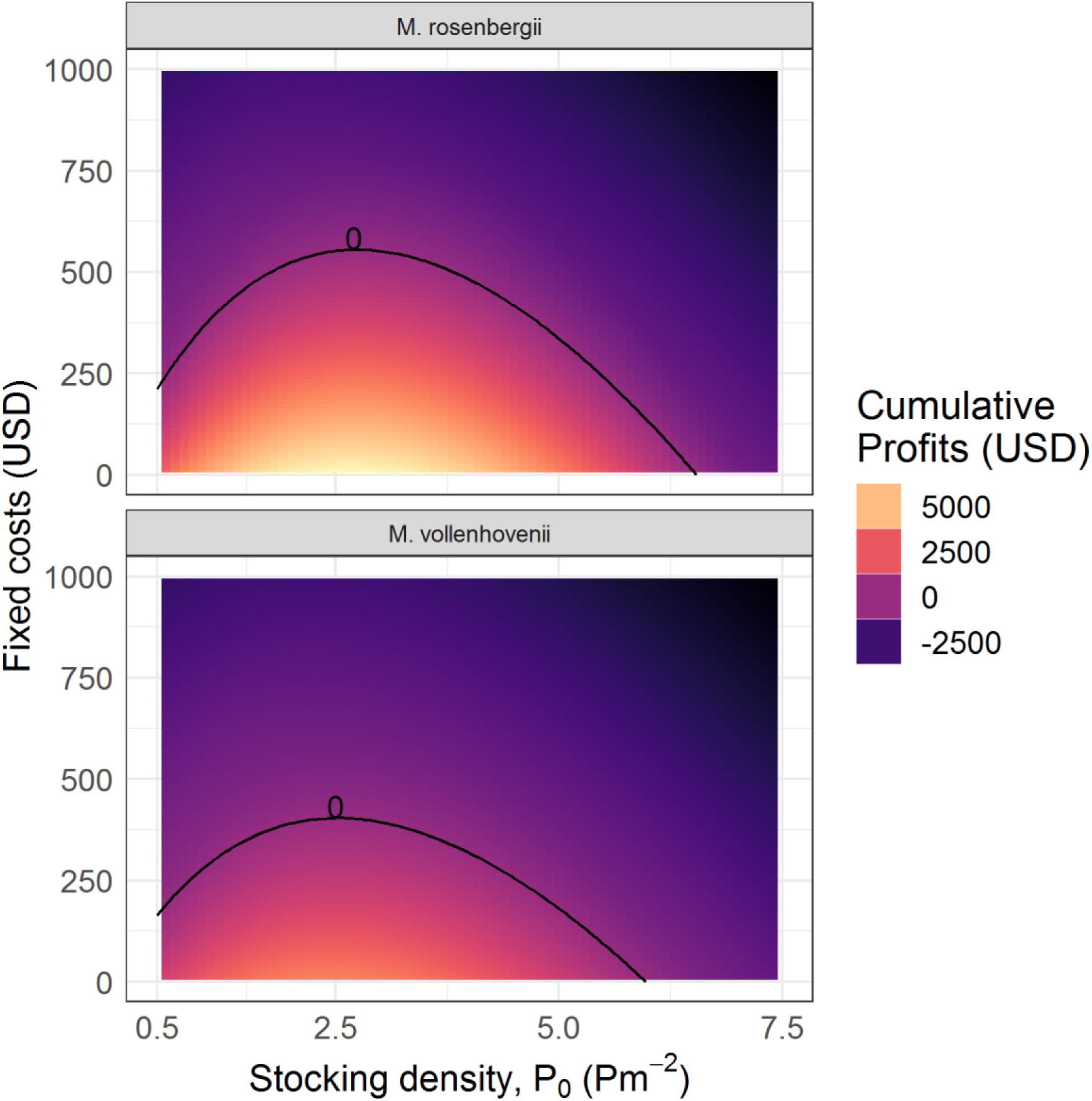
Sensitivity analysis investigating the influence of fixed costs incurred each aquaculture cycle on profits generated by the prawn aquaculture model. Color indicates the 10-year cumulative profits generated by optimal management with stocking at the density indicated on the x-axis. The black line delineates the maximum boundary of profitability.

**Figure S2:**
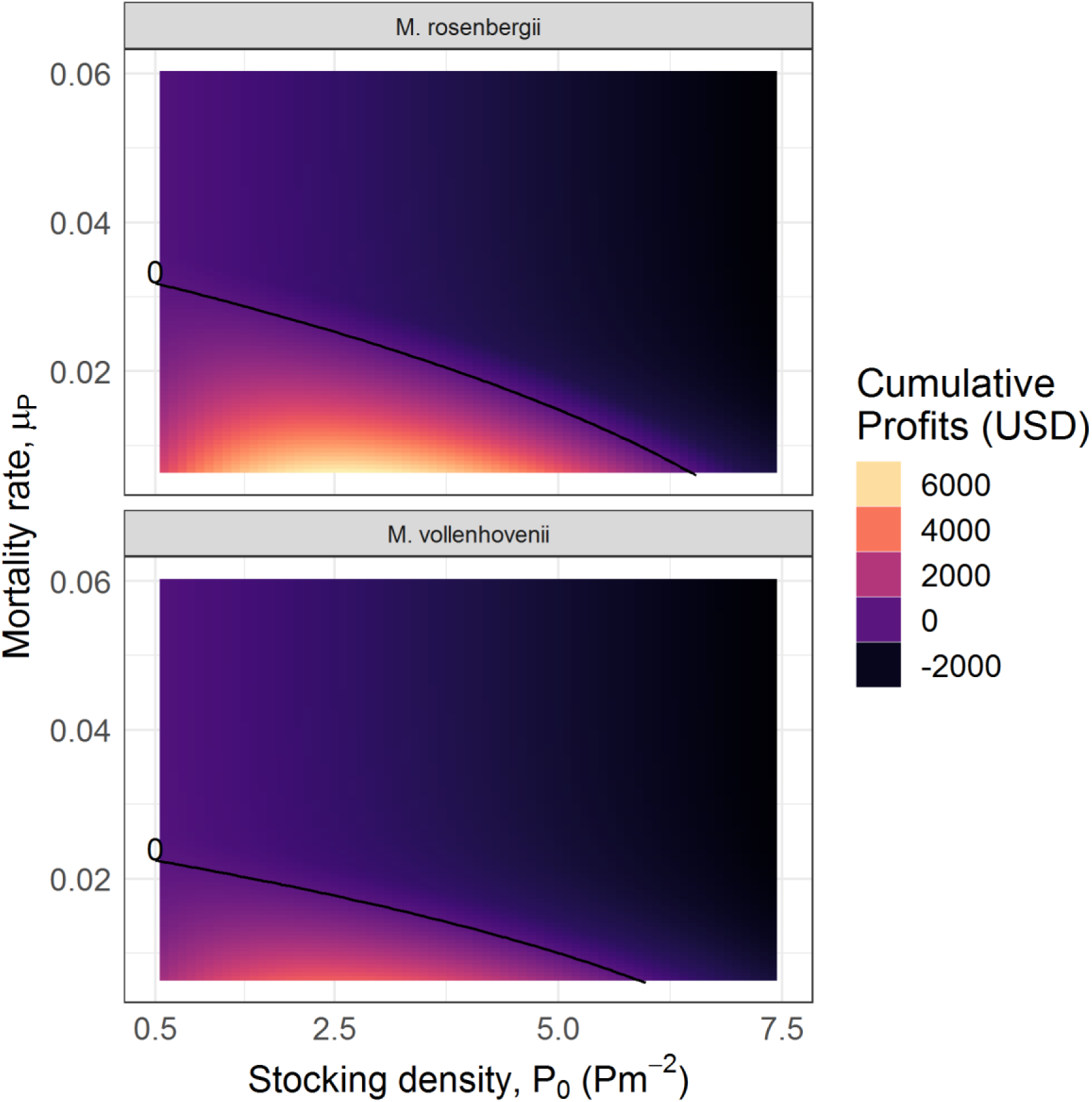
Sensitivity analysis investigating the influence of increased mortality rates (baseline *μ_P_* = 0.006) on profits generated by the prawn aquaculture model. Color indicates the 10-year cumulative profits generated by optimal management with stocking at the density indicated on the x-axis. The black line delineates the maximum boundary of profitability.

**Figure S3:**
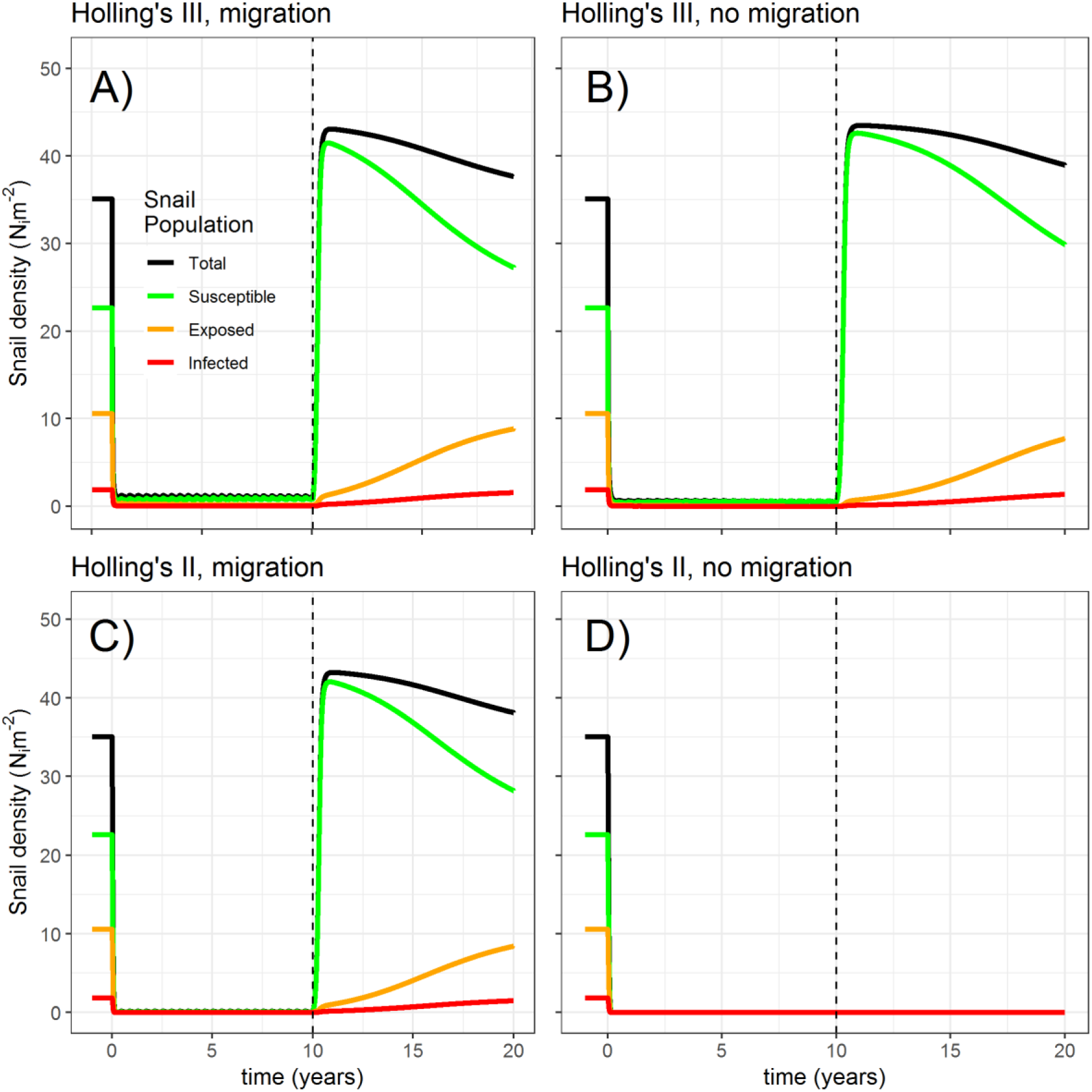
Sensitivity analysis investigating the influence of the predation functional response and migration on snail infection dynamics. The model is simulated assuming a type III functional response and migration from an external source (A), a type III functional response with no migration (B), a type II functional response with migration (C), and a type II functional response with no migration (D). Simulations indicate that as long as migration is present or predation dynamics follow a type III response (indicating decreased predation rates at low densities due to e.g. prey switching or the presence of prey refugia) the snail population rapidly reestablishes after prawn interventions are discontinued.

**Figure S4:**
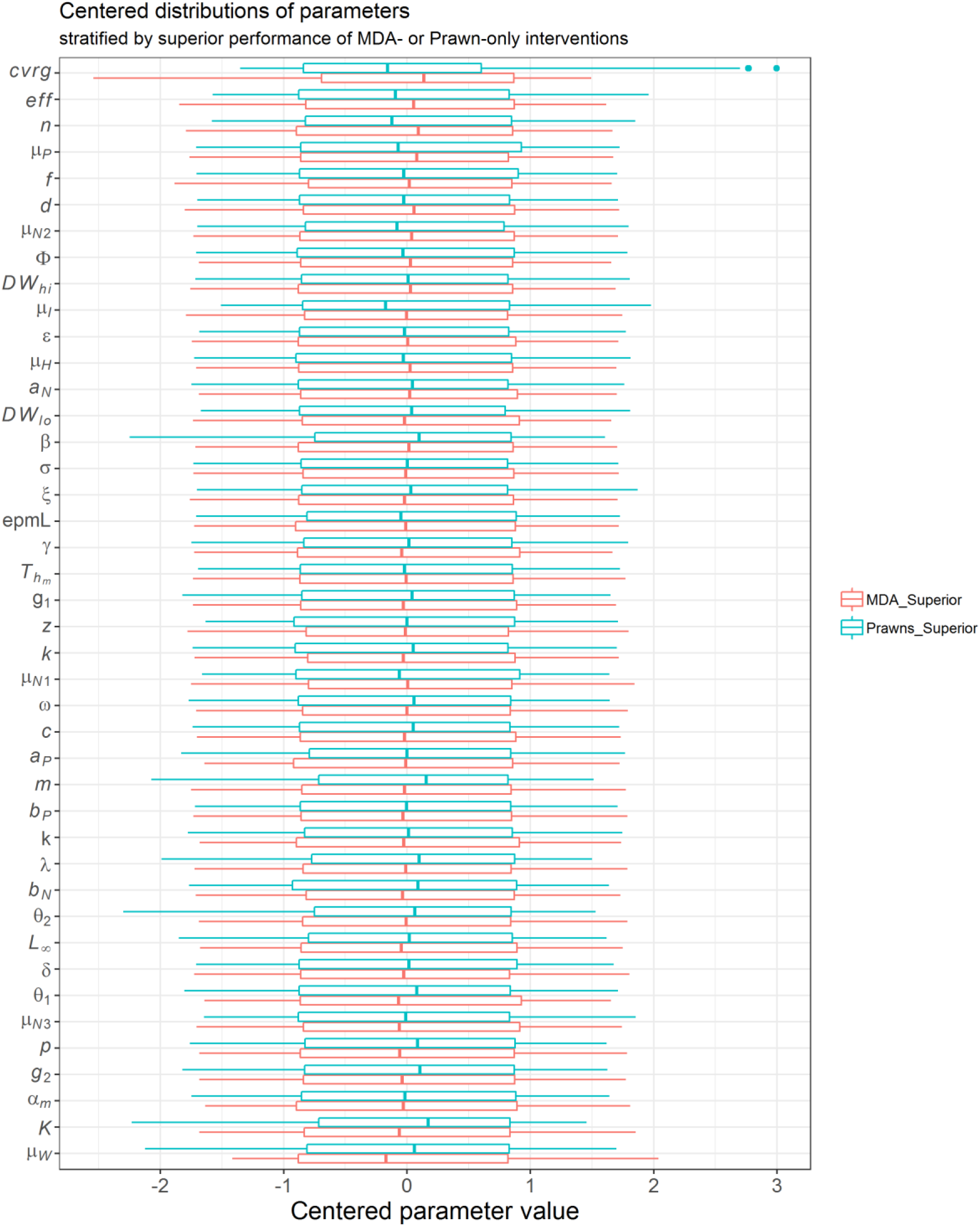
Centered distributions of model parameters stratified by whether more DALYs were averted in MDA-only or in Prawn-only intervention simulations compared to no intervention simulations. Comparisons show that the MDA intervention tends to perform superior when MDA coverage (*cvrg*) and efficacy (*eff*) are higher and when infected snail and prawn mortality are higher (*μ_I_* and *μ_P_*, respectively). The prawn intervention performs notably better when the snail population carrying capacity (K) is higher.

**Figure S5:**
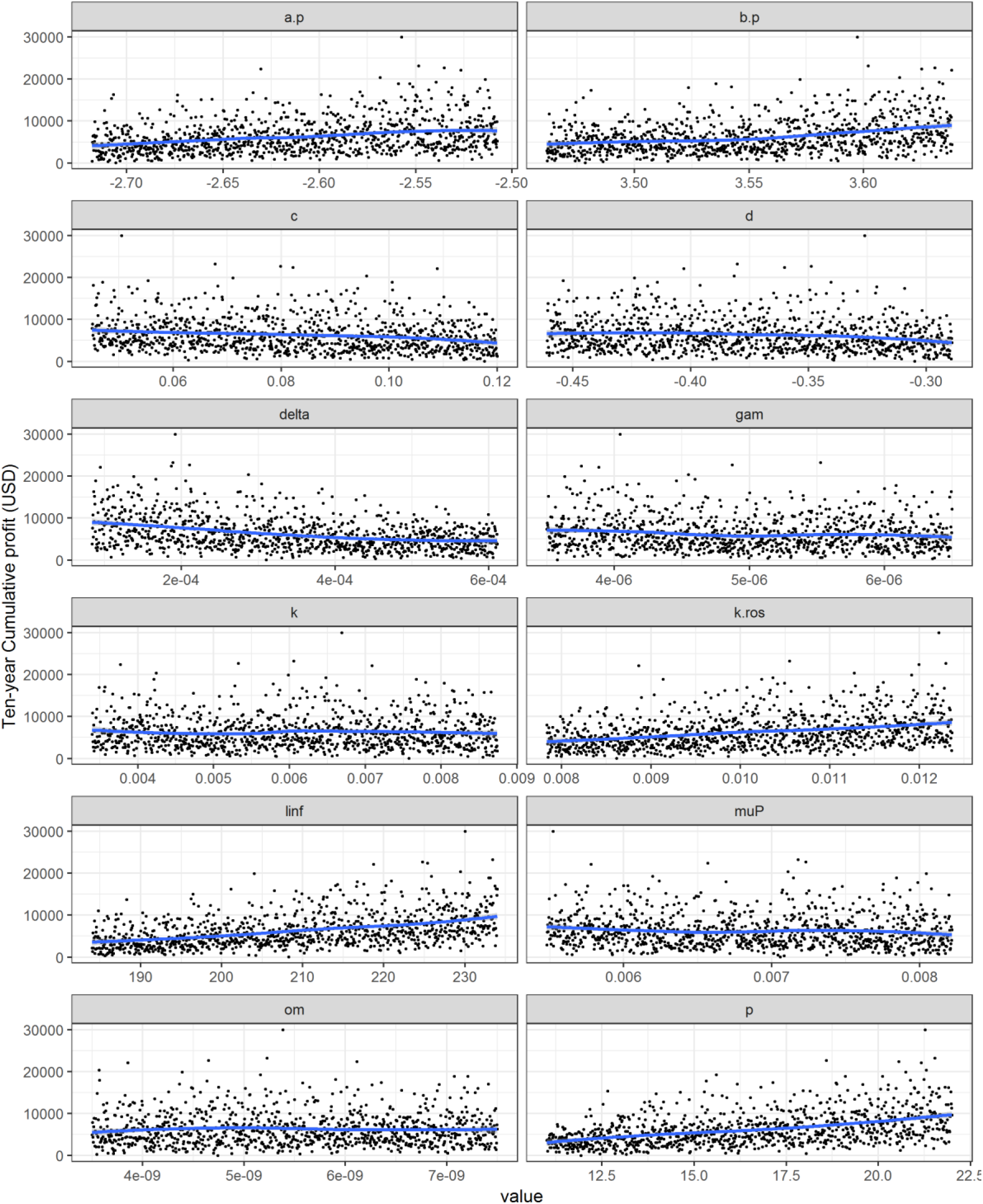
Scatterplots used to verify monotonicity between parameters of the prawn aquaculture model and estimates of ten year cumulative profits. Blue lines represent loess smoothing.

**Figure S6:**
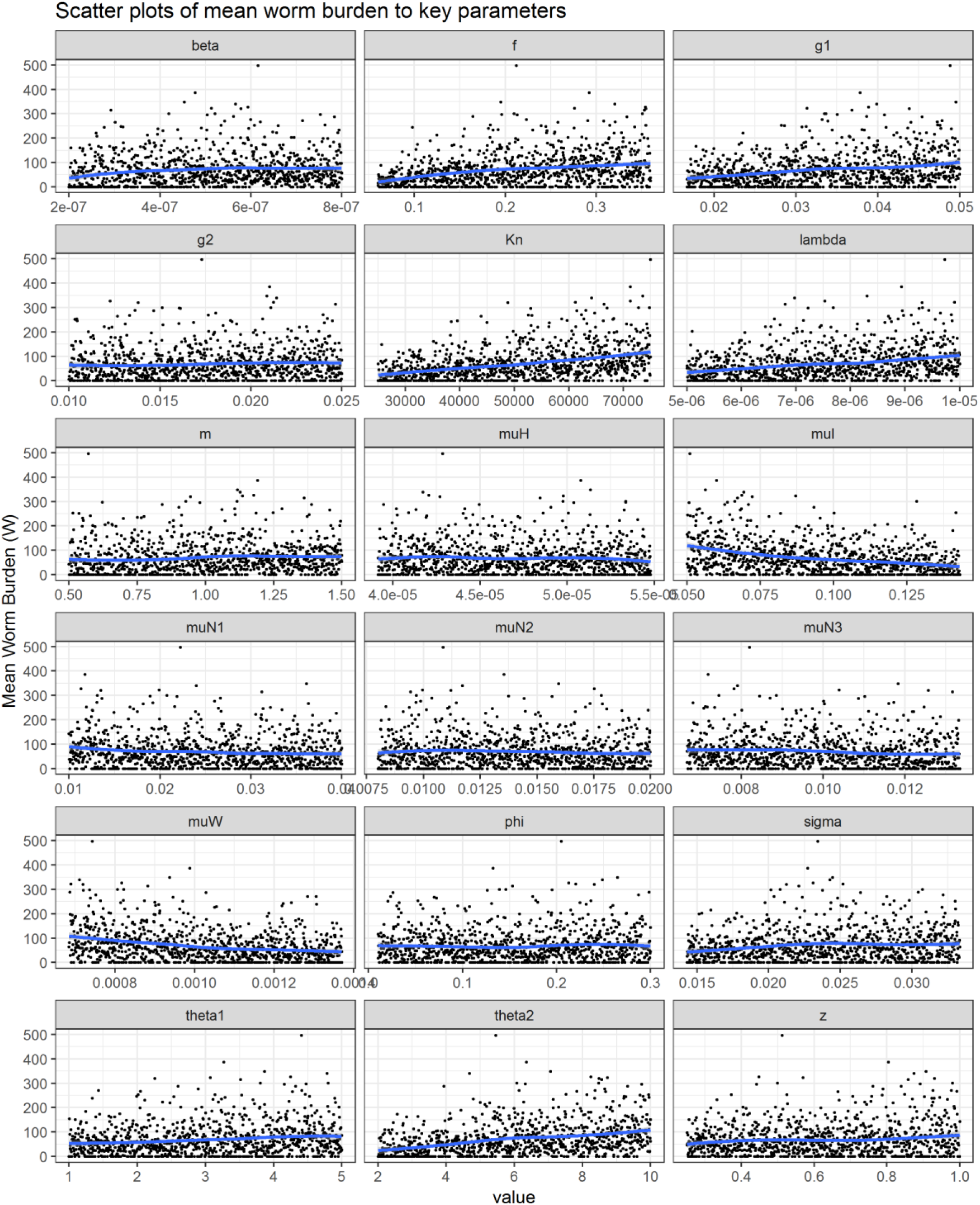
Scatterplots used to verify monotonicity between parameters of the epidemiologic model and estimates of equilibrium mean worm burden. Blue lines represent loess smoothing.

**Figure S7:**
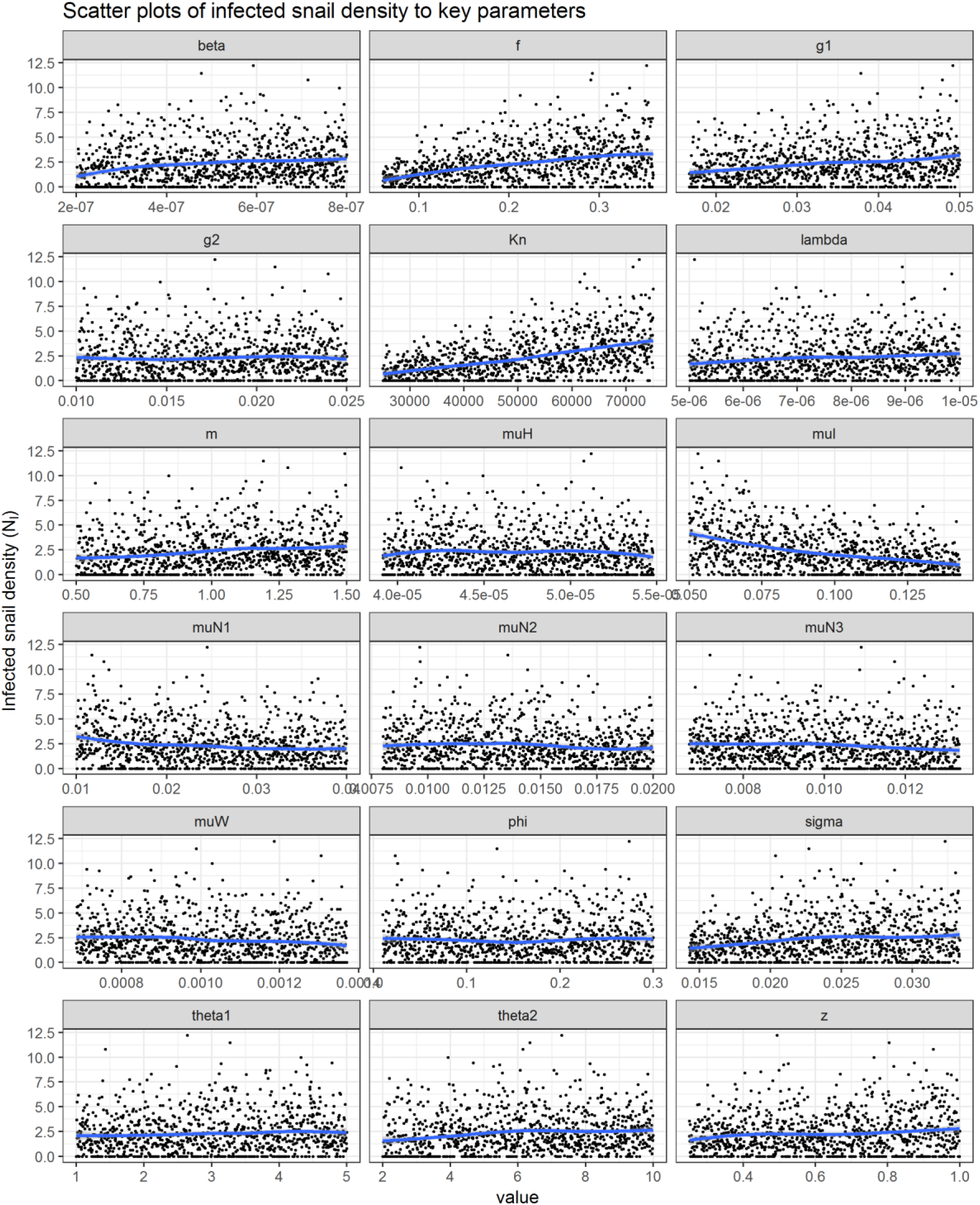
Scatterplots used to verify monotonicity between parameters of the epidemiologic model and estimates of equilibrium infected snail density. Blue lines represent loess smoothing.

**Figure S8:**
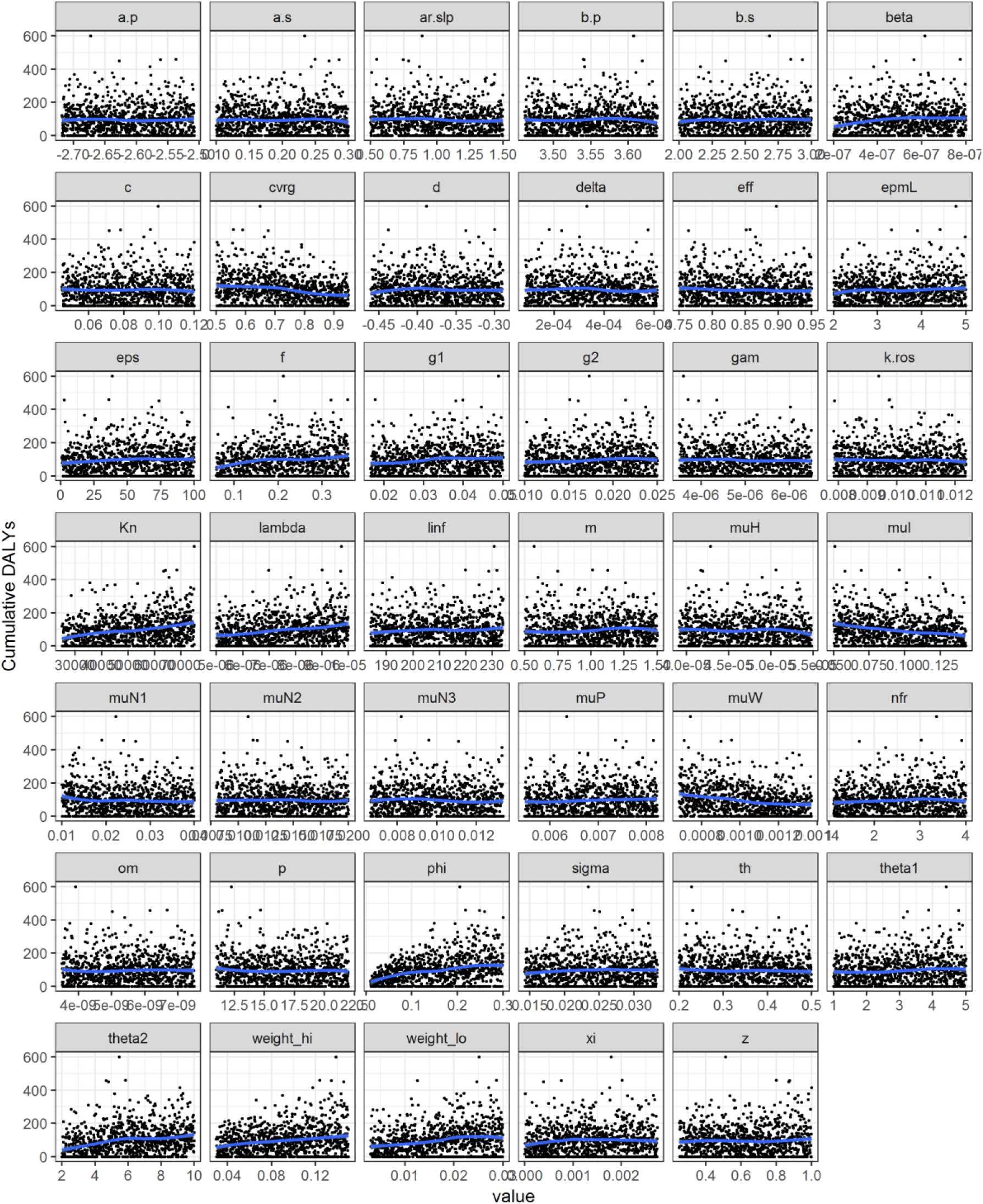
Scatterplots used to verify monotonicity between parameters of the combined model and estimates of DALYs lost during ten years of integrated MDA and prawn intervention. Blue lines represent loess smoothing.

